# Hsf1 phosphorylation generates cell-to-cell variation in Hsp90 levels and promotes phenotypic plasticity

**DOI:** 10.1101/185934

**Authors:** Xu Zheng, Ali Beyzavi, Joanna Krakowiak, Nikit Patel, Ahmad S. Khalil, David Pincus

**Author notes:** Present address: Koch Institute for Integrative Cancer Research, Cambridge, USA. These authors contributed equally (Xu Zheng, Ali Beyzavi).

## Abstract

Clonal populations of cells exhibit cell-to-cell variation in the transcription of individual genes. In addition to this “noise” in gene expression, heterogeneity in the proteome and the proteostasis network expands the phenotypic diversity of a population. Heat shock transcription factor (Hsf1) regulates chaperone gene expression, thereby coupling transcriptional noise to proteostasis. Here we show that cell-to-cell variation in Hsf1 activity is an important determinant of phenotypic plasticity. Budding yeast cells with high Hsf1 activity were enriched for the ability to acquire resistance to an antifungal drug, and this enrichment depended on Hsp90 – a known “phenotypic capacitor” and canonical Hsf1 target. We show that Hsf1 phosphorylation promotes cell-to-cell variation, and this variation – rather than absolute Hsf1 activity – promotes antifungal resistance. We propose that Hsf1 phosphorylation enables differential tuning of the proteostasis network in individual cells, allowing populations to access a wide range of phenotypic states.

## INTRODUCTION

Genetically identical cells grown together in the same environment nonetheless display cell-to-cell variation in gene expression (Colman-Lerner et al., 2005; Elowitz et al., 2002; Raser and O'Shea, 2004, 2005; Weinberger et al., 2005). While most frequently observed in microorganisms, such as bacteria and yeast, gene expression variation is also found in developing mammalian cells and human embryonic stem cells (Silva and Smith, 2008; Stelzer et al., 2015). Such variation has been proposed to be the mechanistic underpinning of lineage commitment during human development, the epithelial to mesenchymal transition in cancer metastasis, organ regeneration in planarians, bacterial persistence in the presence of antibiotics, and the ability of yeast cells to remain fit in fluctuating environments (Harms et al., 2016; Newman et al., 2006; Oderberg et al., 2017; Silva and Smith, 2008; Ye and Weinberg, 2015). While differences in cell size, cell cycle position and chromatin state can partially account for cell-to-cell variation, much of the variability has been attributed to the inherently stochastic process of gene expression (Colman-Lerner et al., 2005; Raj and van Oudenaarden, 2008; Raser and O'Shea, 2005). Despite the underlying stochasticity, gene expression variation itself varies widely across the genome, with some sets of genes showing very low variation among cells (e.g., ribosomal protein genes) and other sets of genes (e.g., stress responsive genes) showing high levels of variation (Newman et al., 2006). Yet, individual genes within these regulons show strong covariance, indicating the source of the variation lies in the activity of upstream transcription factors and signaling pathways (Stewart-Ornstein et al., 2012). As such, cell-to-cell variation may be a property that is under genetic control and can be tuned up and down over evolution.

On top of this gene expression variation, cell-to-cell differences also exist in the state of the proteome. Perhaps the most striking examples of proteome variation come from prion proteins, which can exist in either soluble or self-templating amyloid conformations (Shorter and Lindquist, 2005). Prions have been shown to have the ability to broadly reshape the proteome by challenging chaperones and other components of the protein homeostasis (proteostasis) machinery and even by globally altering protein translation (Serio and Lindquist, 1999; Shorter and Lindquist, 2008). Moreover, chaperones themselves can exist in large heterotypic complexes that differ between cells in what has been termed the “epichaperome,” giving rise to altered susceptibility of cancer cells to drugs that target the essential chaperone Hsp90 (Rodina et al., 2016). By buffering the proteome and stabilizing near-native protein folds, Hsp90 has been shown to mask latent genetic variation in fruit flies and plants and to enhance the ability of yeast cells to acquire novel phenotypes such as resistance to anti-fungal drugs (Cowen and Lindquist, 2005; Queitsch et al., 2002; Rutherford and Lindquist, 1998). In this regard, Hsp90 has been termed a “phenotypic capacitor” (Sangster et al., 2004).

Heat shock factor 1 (Hsf1) regulates the expression of many components of the proteostasis machinery – including Hsp90 – in eukaryotes from yeast to humans (Anckar and Sistonen, 2011). In unstressed budding yeast cells, a different chaperone, Hsp70, binds to Hsf1 and restrains its activity. Upon heat shock, Hsp70 dissociates from Hsf1 leaving Hsf1 free to induce expression of its target genes (Zheng et al., 2016). Heat shock also triggers Hsf1 hyper-phosphorylation. Although phosphorylation is a conserved hallmark of Hsf1 activation, it is dispensable for acute Hsf1 activity during heat shock. Rather than switching Hsf1 on, phosphorylation enables Hsf1 to sustain increased activity during prolonged exposure to elevated temperature (Zheng et al., 2016). Here we identify a novel role for Hsf1 – and Hsf1 phosphorylation – that may have provided a strong selective advantage during evolution. We show that Hsf1 generates cell-to-cell variation in Hsp90 levels, which in turn contributes to the ability of *Saccharomyces cerevisiae* to acquire resistance to the antifungal drug fluconazole. We find that the ability of Hsf1 to become phosphorylated is a key factor in generating population level heterogeneity in its activity. We propose that by coordinately controlling cytosolic chaperone genes including Hsp90, Hsf1 promotes phenotypic plasticity.

## RESULTS

### Differential cell-to-cell variation in Hsf1 activity in response to heat shock and AZC

In addition to heat shock, Hsf1 is known to respond to a variety of chemical stressors that impair proteostasis. In particular, the small molecule azetidine 2-carboxylic acid (AZC) is known to strongly activate Hsf1 (Trotter et al., 2002). AZC is a proline analog that is charged onto tRNA^pro^ and incorporated into nascent proteins during translation, impairing their subsequent folding (Fowden and Richmond, 1963). Our previous mass spectrometry data indicated that Hsf1 displays distinct phosphorylation patterns in cells that had been heat shocked compared to cells treated with AZC (Zheng et al., 2016). To explore these differences, we monitored phosphorylation of FLAG-tagged Hsf1 following heat shock and treatment with AZC by observing its electrophoretic mobility over time by western blot. Heat shock induced progressive and sustained Hsf1 phosphorylation leading to a dramatic shift in its mobility over the heat shock time course (Figure 1A). By contrast, AZC led to only a modest shift in Hsf1 mobility (Figure 1A). Despite these differences in phosphorylation, both heat shock and AZC robustly induced Hsf1 transcriptional activity as measured by flow cytometry of cells expressing HSE-YFP, a fluorescent reporter of Hsf1 transcriptional activity (Figure 1B) (Zheng et al., 2016). While heat shock resulted in rapid HSE-YFP induction, plateauing after 1 hour, AZC induced Hsf1 activity with delayed but sustained kinetics, ultimately leading to the same maximal output level as heat shock after 4 hours (Figure 1B).

**Figure 1.**
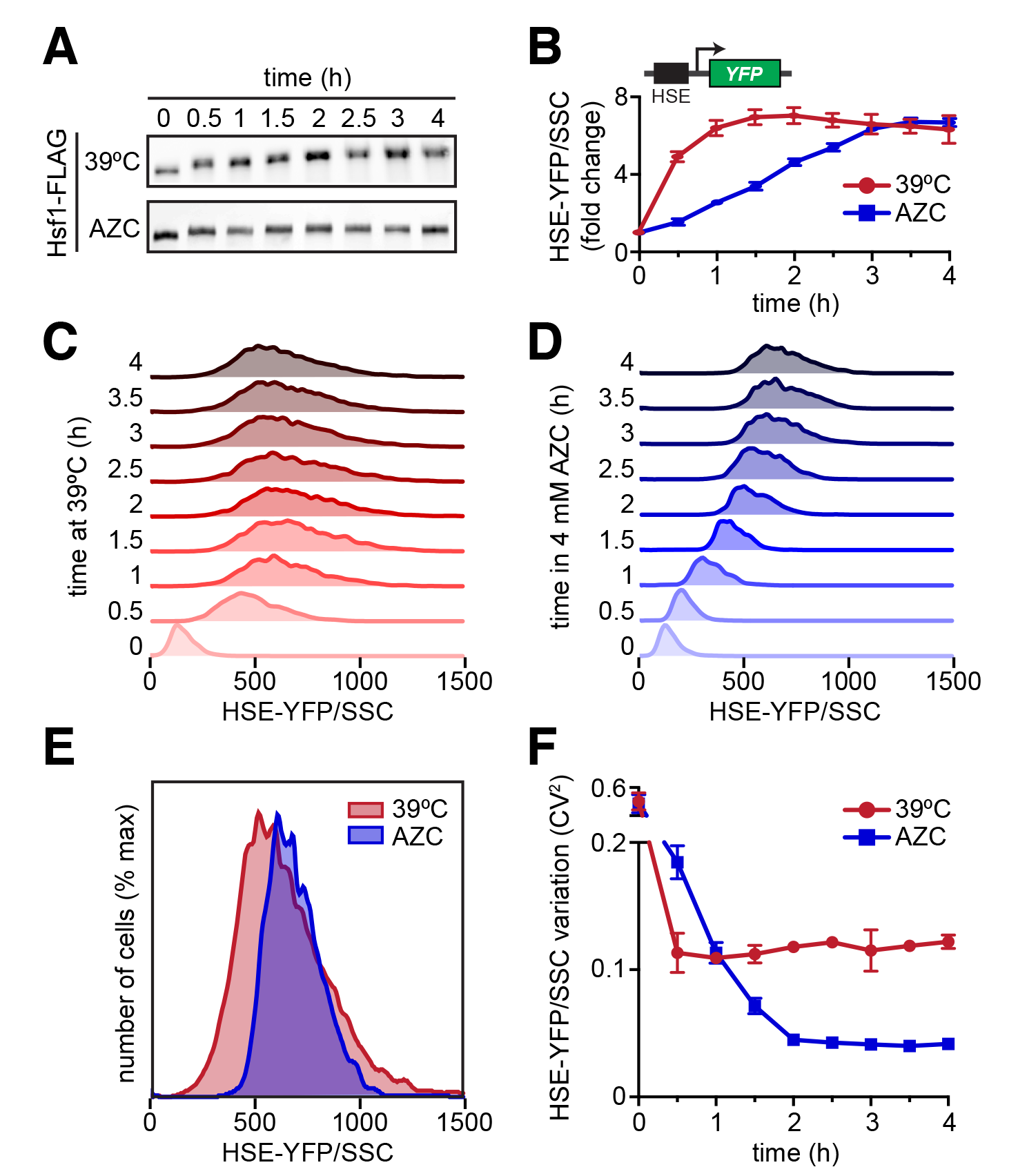
Differential cell-to-cell variation in Hsf1 activity in response to heat shock and AZC. **(A)** Anti-FLAG Western blot showing Hsf1 phosphorylation by electrophoretic mobility shift over time in response to heat shock at 39°C or 4 mM AZC. **(B)** Hsf1 activity quantified by the levels of the HSE-YFP transcriptional reporter measured by flow cytometry over time in response to heat shock at 39°C or 4 mM AZC. The reporter consists of four repeats of the heat shock element (HSE) recognized by Hsf1 upstream of a crippled *CYC1* promoter. For each cell, the YFP level was normalized by side scatter (HSE-YFP/SSC) to control for cell size. Each point represents the median of the fluorescence distribution of 10000 cells averaged over 3 biological replicates; error bars show the standard deviation of the replicates. **(C)** Single cell fluorescence distributions over a 4-hour heat shock time course. For each cell, the YFP level was normalized by side scatter (HSE-YFP/SSC) to control for cell size. **(D)** As in (C), but over a time course of treatment with 4 mM AZC. **(E)** Overlay of the 4-hour time points from (C) and (D). **(F)** Quantification of cell-to-cell variation in HSE-YFP/SSC levels over time courses in (C) and (D) using the square of the coefficient of variation (CV^2^) as a metric.

Along with altered activation kinetics, we observed differences in the single cell fluorescence distributions: the populations of cells that had been heat shocked showed a broad distribution of HSE-YFP levels, while cells treated with AZC showed narrower distributions, indicating reduced levels of cell-to-cell variation in Hsf1 activity. To account for potential differences in cell size, we normalized each cell’s HSE-YFP fluorescence by its size as measured by side scatter (SSC) and plotted the resulting normalized distributions over the heat shock and AZC time courses (Figure 1C, D) (Stewart-Ornstein et al., 2012). Direct comparison of the HSE-YFP/SSC distributions following 4 hours of heat shock or treatment with AZC reveals that despite a slightly higher average level of Hsf1 activation in the AZC-treated cells, cellular heterogeneity is greatly reduced compared to heat shocked cells (Figure 1E). To quantify this cell-to-cell variation in the HSE-YFP reporter, we calculated the square of the coefficient of variation (CV^2^) (Stewart-Ornstein et al., 2012). While the CV^2^ drops as the mean increases for both heat shocked and AZC-treated cells, it is 3-fold higher in heat-shocked cells by the end of the experiment (Figure 1F). These data show that while heat shock and AZC both potently activate Hsf1, they do so with distinctive features. Heat shock triggers rapid Hsf1 activation coupled to high levels of phosphorylation and results in a high degree of cell-to-cell variation in the HSE-YFP reporter. By contrast, AZC induces slow, sustained Hsf1 activation, modest Hsf1 phosphorylation and reduced noise in the HSE-YFP reporter.

### Hsf1 phosphorylation generates cell-to-cell variation during heat shock

Since increased Hsf1 phosphorylation was associated with increased cell-to-cell variation in the HSE-YFP reporter, we wondered if Hsf1 phosphorylation was responsible for increasing noise during heat shock. To test this, we leveraged a non-phosphorylatable mutant we had previously generated, Hsf1^Δpo4^, in which we mutated all 152 possible sites of phosphorylation to alanine (Zheng et al., 2016). We performed a heat shock time course and measured the HSE-YFP reporter by flow cytometry in wild type cells and cells expressing Hsf1^Δpo4^. Indeed, after normalizing for cell size, we observed reduced noise in the HSE-YFP reporter in the Hsf1^Δpo4^ cells compared to wild type throughout the heat shock time course (Figure 2A, B). As an orthogonal approach to measure cell-to-cell variation in Hsf1 activity, we developed a microfluidic-based assay to measure the HSE-YFP reporter in single cells over time in response to heat shock using a novel platform that enables precision temperature control (Figure 2C, Figure S1). In agreement with the flow cytometry data, cells expressing Hsf1^Δpo4^ showed lower noise in HSE-YFP levels over the heat shock time course (Figure 2D, E). Thus, Hsf1 phosphorylation promotes cell-to-cell variation in Hsf1 activity during heat shock.

**Figure 2.**
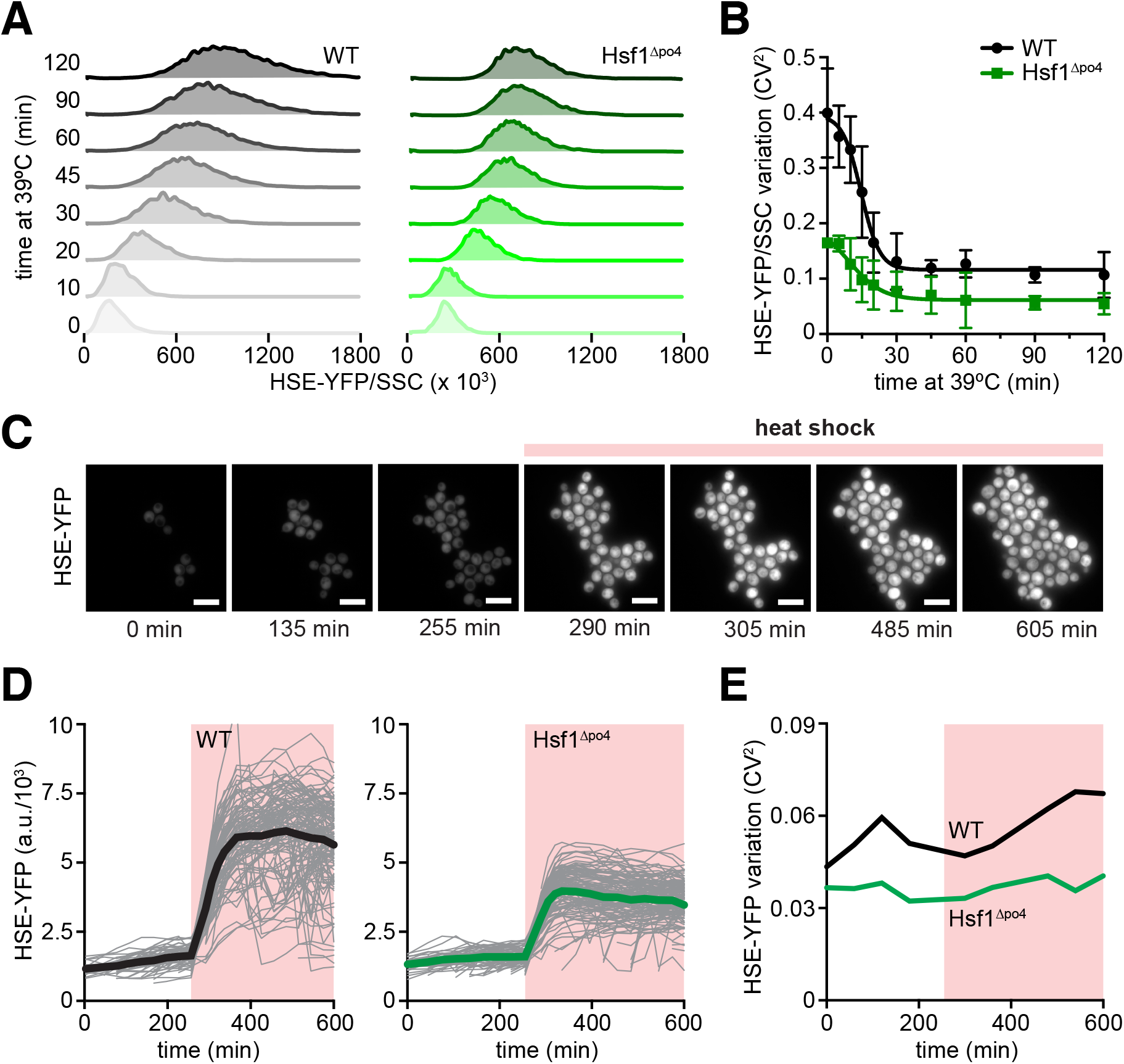
Hsf1 phosphorylation generates cell-to-cell variation during heat shock. **(A)** Single cell fluorescence distributions over a heat shock time course in wild type cells and cells expressing Hsf1^Δpo4^ as the only copy of Hsf1. For each cell, the YFP level was normalized by side scatter (HSE-YFP/SSC) to control for cell size **(B)** Quantification of cell-to-cell variation in HSE-YFP/SSC levels over time courses in (A). **(C)** Images of wild type cells expressing the HSE-YFP reporter growing over time in a microfluidic device at 25°C and shifted to 39°C at the indicated time. Scale bar is 10 μm. **(D)** Quantification of HSE-YFP levels over time in the microfluidic device in wild type cells and cells expressing Hsf1^Δpo4^ as the only copy of Hsf1. For each cell, the total YFP fluorescence was divided by cell area to account for cell size differences. Each gray trace represents the trajectory of a single cell and the thick lines show population averages **(E)** Quantification of cell-to-cell variation in HSE-YFP levels over time courses in (D).

### Hsp90 expression displays Hsfl-dependent cell-to-cell variation

To investigate functional roles for cell-to-cell variation in Hsf1 activity, we monitored expression of Hsp90, an endogenous Hsf1 target gene known to influence phenotypic plasticity (Cowen and Lindquist, 2005). Budding yeast encodes two Hsp90 paralogs, Hsc82 and Hsp82, with Hsc82 being highly expressed under all conditions and Hsp82 showing both basal and inducible expression in response to heat shock (Solis et al., 2016). We tagged Hsp82 with YFP and monitored its expression in single cells expressing either wild type Hsf1 or Hsf1^Δpo4^ by fluorescence microscopy and flow cytometry. Consistent with the HSE-YFP reporter, Hsp82-YFP expression showed greater variability among wild type cells than Hsf1^Δpo4^ cells (Figure 3A, B).

**Figure 3.**
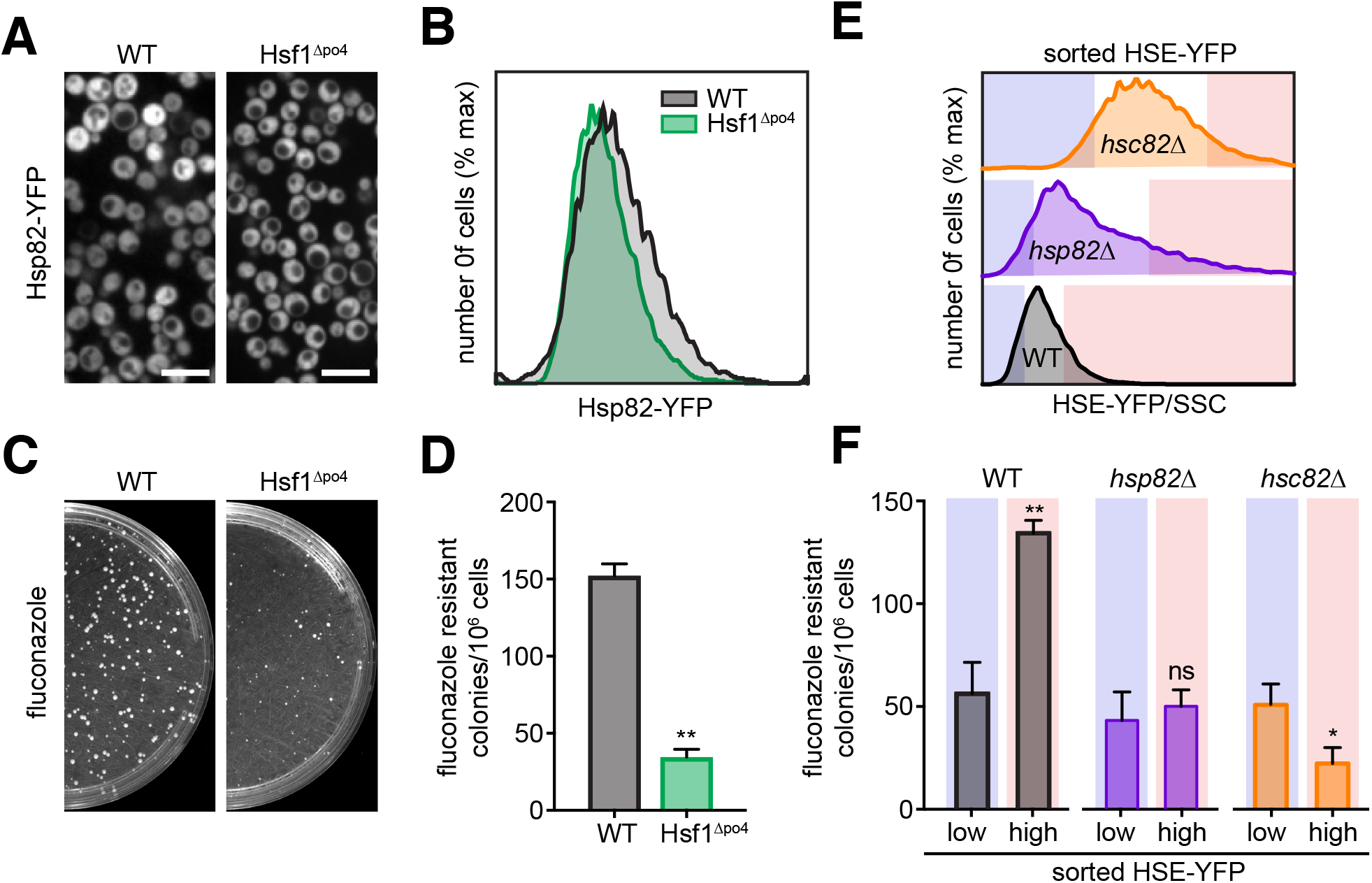
Cell-to-cell variation in Hsf1 activity promotes antifungal resistance in an Hsp90-dependent manner. **(A)** Spinning disc confocal images of wild type cells and Hsf1^Δpo4^ cells expressing Hsp82-YFP. Scale bar is 10 μm. **(B)** Single cell fluorescence distributions of wild type cells and Hsf1^Δpo4^ cells expressing Hsp82-YFP as determined by flow cytometry For each cell, the YFP level was normalized by side scatter (Hsp82-YFP/SSC) to control for cell size **(C)** Appearance of fluconazole-resistant colonies in wild type cells and Hsf1^Δpo4^ cells. 10^6^ cells were plated on YPD plates supplemented with 128 μg/ml fluconazole and incubated at room temperature for 4 days. **(D)** Quantification of the number of fluconazole resistant colonies from three biological replicates of the experiment shown in (C). Error bars show the standard deviation of the replicates (* p < 0.05; ** p < 0.01 by two-tailed T-test) **(E)** Single cell fluorescence distributions of HSE-YFP/SSC in wild type and Hsp90 mutant cells. under basal conditions. Deletion of *HSP82* leads to a modest increase in HSE-YFP levels, while deletion of *HSC82* leads to a pronounced increase in HSE-YFP levels. The blue and red boxes indicate the tails of the distribution with the highest and lowest expressing 10% of cells. **(F)** Quantification of the number of fluconazole resistant colonies in the tails of the distributions shown in (E) (* p < 0.05; ** p < 0.01 by two-tailed T-test)

Prior studies have implicated Hsp90 as a “phenotypic capacitor” that promotes the ability of cells to acquire novel phenotypes (Sangster et al., 2004). In particular, in fungi, Hsp90 has been shown to increase the rate at which cells acquire resistance to the anti-fungal drug fluconazole (Cowen and Lindquist, 2005). We hypothesized that the increased cell-to-cell variation in Hsp82-YFP expression levels in wild type cells compared to Hsf1^Δpo4^ cells would translate into increased fluconazole resistance. Indeed, wild type cells generated significantly more fluconazole resistant colonies than Hsf1^Δpo4^ cells (Figure 3C, D).

### Acquired fluconazole resistance is enriched in cells with high Hsf1 activity

Since noise in Hsp82-YFP levels and the HSE-YFP reporter correlated with phenotypic plasticity, we wondered if individual cells with elevated HSE-YFP levels would be more likely to become resistant to fluconazole. To test this, we used fluorescence activated cell sorting (FACS) to isolate the tails of the HSE-YFP distribution – the 10% of cells with the highest and lowest HSE-YFP expression levels – in wild type cells, *hsp82Δ* and *hsc82Δ* cells (Figure 3E). In wild type cells, the tail with high HSE-YFP expression was enriched for fluconazole resistant colonies over the low expression tail (Figure 3F). By contrast, the high expression tails of the *hsp82Δ* and *hsc82Δ* cells showed no enrichment, with the *hsc82Δ* high expression tail actually showing decreased resistance (Figure 3F). These results indicate that cells with higher levels of Hsf1 activity have an increased ability to acquire resistance to an antifungal drug, and this ability depends on Hsp90.

### Fluconazole resistance correlates with variation in Hsf1 activity, not its magnitude

Since the high expression tail of the HSE-YFP distribution was enriched for cells able to acquire resistance to fluconazole, we hypothesized that if we could increase the average HSE-YFP expression, we could increase the ability of cells to acquire fluconazole resistance. To test this, we expressed wild type Hsf1 from a promoter under the control of a synthetic, estradiol-responsive (ER) transcription factor (Zheng et al., 2016). In this way, we could titrate the amount of estradiol to tune the expression level of Hsf1, and thereby control the expression level of its target genes and the HSE-YFP reporter. In parallel, we also expressed Hsf1^Δpo4^ using the same system to determine if the deficit in cell-to-cell variation could be overcome by increasing the absolute expression (Figure 4A). We grew the wild type and Hsf1^Δpo4^ cells in a dilution series of estradiol, maintained growth at OD_600_ < 0.5 for 18 hours and measured the HSE-YFP reporter. Estradiol induced dose-dependent induction of the HSE-YFP in both wild type and Hsf1^Δpo4^ cells, though with a steeper curve for Hsf1^Δpo4^ cells (Figure 4B, C). At all doses of estradiol, wild type had higher CV^2^ in the HSE-YFP reporter than Hsf1^Δpo4^ cells (Figure 4B, D). We tested the ability of cells expressing wild type Hsf1 and Hsf1^Δpo4^ at low (1 nM), medium (8 nM) and high (32 nM estradiol) levels to acquire resistance to fluconazole (Figure 4E). Surprisingly, increasing median HSE-YFP did not increase resistance: both wild type and Hsf1^Δpo4^ cells showed maximal resistance at low Hsf1 expression (Figure 4F). However, wild type cells showed greater resistance at all Hsf1 expression levels than Hsf1^Δpo4^ cells (Figure 4F). Thus, increased variation in HSE-YFP levels – not increased median levels – positively correlates with fluconazole resistance (Figure 4G-I).

**Figure 4.**
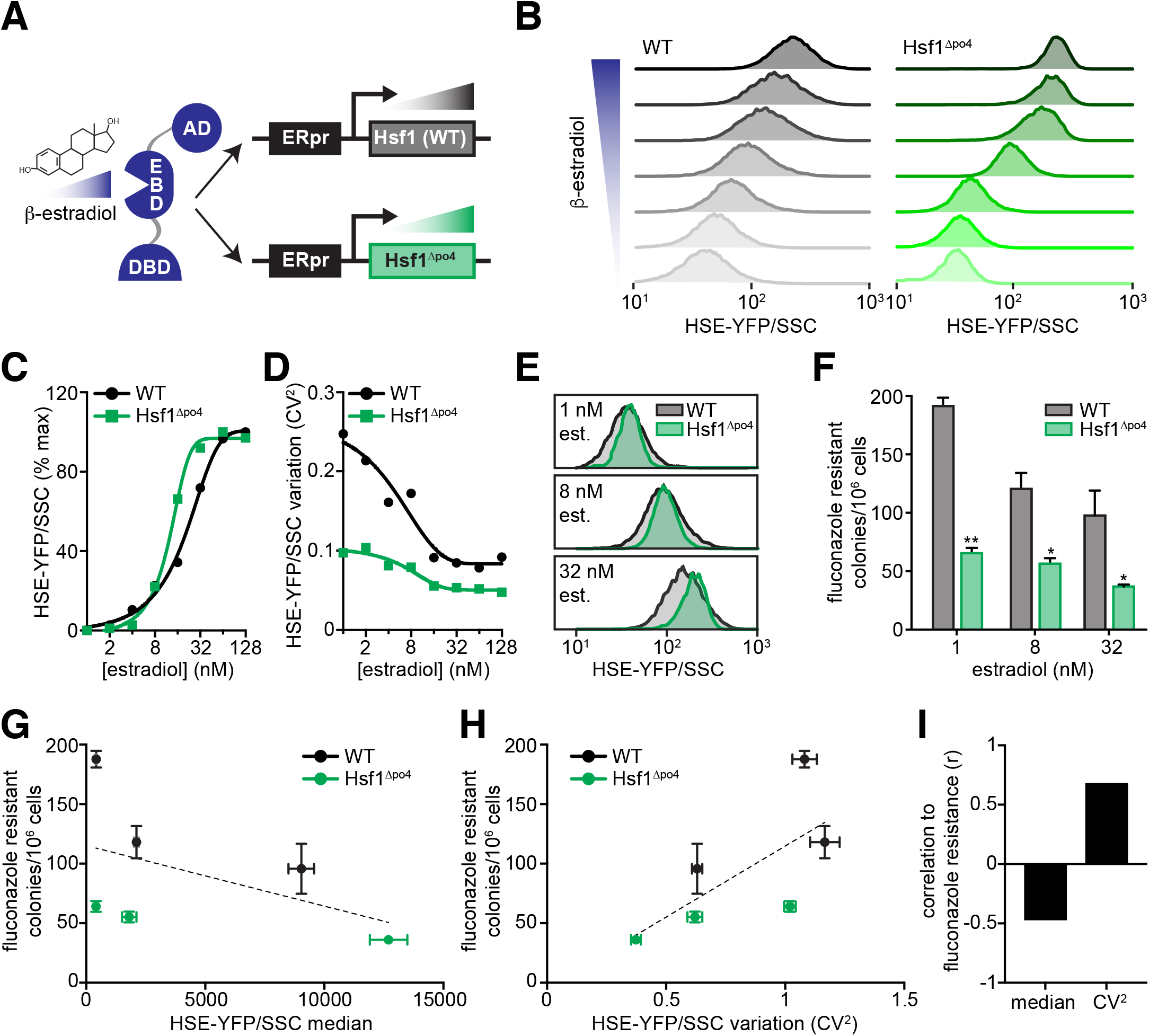
Fluconazole resistance correlates with cell-to-cell variation in Hsf1 activity. **(A)** Schematic showing the estradiol-inducible system used to titrate expression of wild type Hsf1 and Hsf1^Δpo4^. A chimeric transcription factor containing the ligand binding domain of the human estrogen receptor drives the expression of Hsf1 as a function of the concentration of estradiol added to the growth media. **(B)** Single cell HSE-YFP/SSC fluorescence distributions in cells treated across a dose response of estradiol, ranging from 2-128 nM in cells expressing wild type Hsf1 (gray) or Hsf1^Δpo4^ (green) as the only copy of Hsf1. Cells were incubated in log phase for 18 hours in the presence of the indicated dose of estradiol to achieve steady state before measuring. **(C)** Quantification of the median HSE-YFP/SSC across the estradiol dose response in wild type and Hsf1^Δpo4^ cells relative to the maximum value. **(D)** Quantification of the cell-to-cell variation (CV^2^) of the HSE-YFP/SSC distributions across the estradiol dose response in wild type and Hsf1^Δpo4^ cells. **(E)** Overlays of the HSE-YFP/SSC distributions from wild type and Hsf1^Δpo4^ cells treated with the indicated concentrations of estradiol. **(F)** Quantification of fluconazole resistant colonies in wild type and Hsf1^Δpo4^ cells in the presence of the indicated doses of estradiol (* p < 0.05; ** p < 0.01 by two-tailed T-test). **(G)** Fluconazole resistant wild type and Hsf1^Δpo4^ colonies plotted as a function of median Hsf1 activity as measured by the HSE-YFP reporter. Dashed line is a linear regression to all six data points showing a negative correlation. **(H)** Fluconazole resistant wild type and Hsf1^Δpo4^ colonies plotted as a function of the variation in Hsf1 activity as measured by the CV^2^ of the HSE-YFP reporter. Dashed line is a linear regression to all six data points showing a positive correlation. **(I)** Correlation coefficients (r) of the data plotted in (G) and (H).

## DISCUSSION

Cell-to-cell variation in gene expression and the state of the proteome have been proposed to contribute to adaptability in populations of genetically identical cells. Here we identify a single regulatory mechanism that couples variation at the transcriptional and proteomic levels: phosphorylation of the transcription factor Hsf1. We found that when wild type Hsf1 is activated and concomitantly hyperphosphorylated – as is the case during heat shock – cells display high variation in their expression of downstream Hsf1 target genes. By contrast, when Hsf1 is activated without hyperphosphorylation following treatment with AZC, cells show more uniform target gene expression. Moreover, removing the ability of Hsf1 to be phosphorylated reduced cell-to-cell variation. Since the Hsf1 regulon consists of chaperones and other proteostasis factors, variation in Hsf1 activity is likely to lead to variation in the proteostasis network and the state of the proteome.

Functionally, we demonstrated that cell-to-cell variation in Hsf1 activity is correlated with the ability of cells to acquire resistance to the antifungal drug fluconazole, in large part via expression of the known phenotypic capacitor Hsp90. While wild type cells with high levels of Hsf1 activity were enriched for the ability to acquire fluconazole resistance, cells with reduced levels of Hsp90 lost this enrichment. This observation suggested that cells with mor
e Hsf1 activity would have more Hsp90 and therefore show increased antifungal resistance. However, synthetically increasing Hsf1 activity resulted in diminished fluconazole resistance. Moreover, increased expression of non-phosphorylatable Hsf1^Δpo4^ – which showed reduced cell-to-cell variation in its activity compared to wild type Hsf1 – could not compensate for its reduced ability to support fluconazole resistance. Thus, antifungal resistance correlates with variation in Hsf1 activity rather than with the magnitude of Hsf1 activity.

How does Hsf1 phosphorylation lead to cell-to-cell variation and why does variation promote fluconazole resistance? It is likely that differential activation of cell cycle, metabolic and stress responsive kinase pathways results in differential Hsf1 phosphorylation states and leads to distinct levels of transcriptional activity in single cells. However, the finding that variation in Hsf1 activity rather than its absolute activity drives resistance is more intriguing. Although cells with synthetically elevated Hsf1 activity produce more Hsp90, they also overproduce the rest of the Hsf1 regulon, which may be maladaptive. In addition, with forced Hsf1 activity, cells are unable to dynamically regulate Hsf1 to precisely tune its activity according to need. While a high level of Hsf1 activity – and thus Hsp90 levels – may be beneficial early in the process of acquiring resistance to fluconazole, it may be that a subsequent reduction in Hsf1 activity promotes proliferation. The capacity for dynamic and coordinated control over the proteostasis network may endow cells with the plasticity required to maintain fitness in fluctuating environments. By enabling both population-level variation and dynamic control over the proteostasis network, Hsf1 is a powerful phenotypic capacitor.

## ACKNOWLEDGEMENTS

We dedicate this paper to Susan Lindquist and her scientific legacy. We are grateful to K. Reynolds and R. Ranganathan for beneficial discussions, to the Whitehead Institute FACS facility for technical assistance and to N. Azubuine and T. Nanchung for a constant supply of plates and media. This work was supported by an NIH Early Independence Award (DP5 OD017941-01 to D.P.) and a National Science Foundation CAREER Award (MCB-1350949 to A.S.K.).

## AUTHOR CONTRIBUTIONS

Conceptualization, D.P. and A.S.K; Methodology, X.Z., A.B. N.P.; Investigation, X.Z., A.B., J.K., D.P., N.P.; Writing, D.P. and A.S.K; Funding Acquisition, D.P. and A.S.K.; Supervision, D.P. and A.S.K.

## METHODS Yeast strains and cell growth

### Yeast cells were cultured in SDC media as described (Zheng et al., 2016)

| DPY number | Description | Genotype |
| --- | --- | --- |
| 144 | WT HSE-YFP | W303 MATa 4xHSE-Venus::LEU |
| 118 | Hsf1-3xFLAG/V5 | W303 MATa *hsf1*Δ::KAN Hsf1-3xFLAG-V5::TRP1 |
| 416 | Hsf1Δpo4 HSE-YFP | W303 MATa *hsf1*Δ::KAN 4xHSE-Venus::LEU2 Hsf1Δpo4::TRP1 |
| 1221 | *hsc82*Δ HSE-YFP | W303 MATa 4xHSE-Venus::LEU2 *hsc82*Δ::HYG |
| 1222 | *hsp82*Δ HSE-YFP | W303 MATa 4xHSE-Venus::LEU2 *hsp82*Δ::HYG |

### Flow cytometry

Flow cytometry was performed as described (Zheng et al., 2016). Data were processed in FlowJo 10. Data were left ungated and YFP fluorescence was normalized by side scatter (SSC) for each cell.

### Microfluidic heat shock time courses

Single cell heat shock experiments were performed with a custom microfluidic device that uses microscale, on-chip heaters to enable programmable thermal perturbations. The multilayer device was fabricated out of the silicone elastomer polydimethylsiloxane (PDMS/Sylgard 184, Dow Corning) using soft lithographic techniques, as described previously (Duffy et al., 1998; Thorsen et al., 2002; Unger et al., 2000; Vega et al., 2012). The device was aligned and sealed to a pre-cleaned No. 1.5 glass coverslip (Fisher Scientific), onto which micro-scale heater and resistance temperature detector (RTD) wires were patterned. Briefly, a thin layer of Ti (100 Angstroms) was deposited on the glass, followed by a layer of Pt (150 nm) and finally a layer of SPR 220-7 positive photoresist (MicroChem Corp.). After transferring the heater/RTD pattern to the photoresist using a high-resolution transparency photomask (CAD/Art Services, Inc.), the Pt/Ti layers were etched in Aqua Regia solution to generate the final serpentine pattern of heater and RTD wires. The photoresist was then washed off with acetone, and a layer of SiO_2_ (300 nm) was deposited on the final pattern. After assembly, the RTD of each device was calibrated and confirmed to respond linearly to changes in the input temperature, as controlled by a hot plate, across the operating temperature range of 25°C to 45°C (Figure S1). Additionally, the devices were characterized by applying user-defined voltages to the heater and measuring on-chip temperatures using a Flir A655sc high-resolution long-wave infrared thermal camera (Flir Systems, Inc.) to validate the device’s ability to achieve accurate and stable heat shock conditions (Figure S1).

To perform microfluidic experiments, we used a custom microfluidic platform that controls the delivery of liquids to the device and the actuation of valves (Vega et al., 2012). The voltage across the heater wire and the resistance across the RTD wire were controlled and measured, respectively, to modulate the on-chip temperature. For each experiment, cells were inoculated 1:100 from overnight SDC cultures into 2 mL SDC and allowed to grow 3-4 hours (OD ~0.5) before seeding the device. Cells were then loaded into the device to trap a small number of cells in growth chambers. Cells were grown on-chip in SDC media for 4-5 hours at 25°C, ambient conditions maintained with a Controlled Environment Microscope Incubator (Nikon Instruments, Inc.) designed for live-cell imaging. Cells were then subjected to a heat shock by applying voltage across the heater wire to ramp to and maintain an on-chip temperature of 39°C.

Throughout, images were collected at 15-minute intervals at 100x magnification (Plan Apo Lambda 100X, NA 1.45) using an Eclipse Ti-E inverted microscope (Nikon Instruments, Inc.) equipped with the “Perfect Focus System”, a XYZ-motorized stage, and a Clara-E charge-coupled device (CCD) camera (Andor Technology). Images were acquired in phase contrast configuration and in the YFP fluorescent channel. Filters and light sources (Nikon LED and Lumencor SPECTRA X Light Engine) were automatically controlled using the supplier’s software (NIS-Elements Advanced Research). Following each experiment, cells were segmented using custom image analysis software written for Matlab (Mathworks, Natick, MA), and fluorescence values for each cell, averaged over cell area, were obtained from the time series of YFP images.

### Spinning disc confocal imaging

Imaging was performed as described (Zheng et al., 2016).

### Fluconazole acquired resistance assay

Wild type and Hsf1^Δpo4^ cells were grown to mid-log phase and 10^6^ cells were plated on YPD plates with 128 μg/ml fluconazole + 25 nM estradiol. Plates were incubated for 4 days at room temperature. Colonies were counted from 3 biological replicates and mean and standard deviation were calculated.

### Fluorescence activated cell sorting

10^6^ cells from the high‐ and low-expressing tails (top and bottom 10% of cells) of the HSEYFP/SSC distribution in wild type, *hsp82Δ* and *hsc82Δ* cells were sorted in the Whitehead Institute FACS facility on a BD FACS Aria.

### Estradiol dose responses

Performed as described (Zheng et al., 2016).

**Figure S1.**
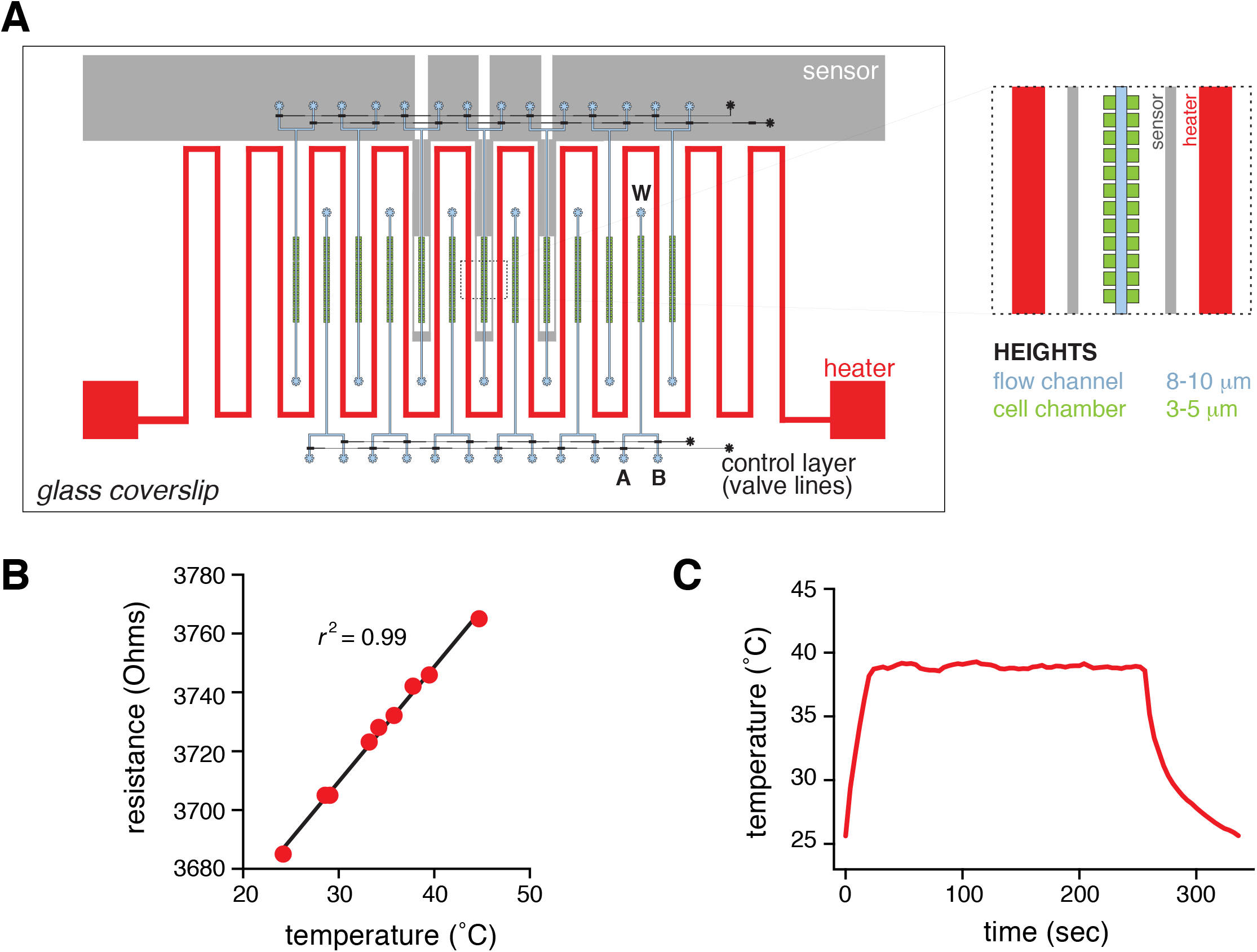
Characterization of the microfluidic heat shock device. **(A)** Schematic of the microfluidic heat shock device. The assembled device consists of a glass coverslip patterned with Pt/Ti heater and sensor wires (grey), and a multilayer PDMS device with flow (black and blue channels) and control (red channels) layers. Cells loaded from either port A or B are trapped in chambers (blue) that have been fabricated to the height of a single monolayer of S. *cerevisiae* cells. W denotes waste port. **(B)** Representative calibration curve for the on-chip sensor showing a linear response across the operating temperature range. **(C)** Using the on-chip heater to achieve and accurately maintain chamber temperature of 39°C, as measured by a high-resolution long-wave infrared thermal camera.

